# *IN SILICO* SCREENING AND MOLECULAR DYNAMIC SIMULATION STUDIES OF POTENTIAL SMALL MOLECULE IMMUNOMODULATORS OF THE KIR2DS2 RECEPTOR

**DOI:** 10.1101/2020.05.10.087148

**Authors:** Adekunle Babajide Rowaiye, Jide Olubiyi, Doofan Bur, Ikemefuna Chijioke Uzochukwu, Alex Akpa, Charles Okechukwu Esimone

## Abstract

The World Health Organization reports that cancer is one of the most common causes of death worldwide and it accounted for an estimated 9.6 million deaths in 2018. As compared with chemotherapy or radiotherapy, immunotherapy offers a safer, less stressful and selective strategy in the destruction of cancer cells. The killer cell immunoglobulin-like receptor 2DS2 (KIR2DS2) expressed on Natural Killer (NK) cells are involved in signal transduction processes that produce pro-inflammatory cytokines and directly destroy cancer and virally infected cells. The aim of this study is to identify small molecules from natural products that have strong binding affinity with KIR2DS2 and possible bioactivity. A library of small molecule natural compounds obtained from edible African plants was used for *in Silico* molecular docking simulations of KIR2DS2 (PDBID: 1m4k) using *Pyrx*. An arbitrary docking score ≥ −7.0 kcal/mol was chosen as cut off value. Screening for drug-likeness and ligand efficiency was based on the molecular descriptors of the compounds as provided by *Pubchem*. Further screening for saturation, molar refractivity, promiscuity, pharmacokinetic properties, and bioactivity was done using *SWISSADME, PKCSM*, and *Molinspiration* respectively. The molecular dynamic simulation and analyses was done using the *Galaxy* webserver which uses the GROMACS software. Analyses of molecular dynamic simulation were done using *Galaxy* and *MDWEB* webservers. Gibberellin A20 and A29 were obtained as the lead compounds and they show better promise as drug candidates for KIR2DS2 than the standard. It is recommended that the immuno-stimulatory effect of the lead compounds on KIR2DS2 be further investigated.

## 1.0 Introduction

The Killer cell immunoglobulin-like receptors (KIRs) are usually located on a sub set of T cells and NK cells. These proteins are coded for by the polymorphic KIR genes, which translate into both activating (aKIR) and inhibiting (iKIR) receptors [1]. The KIR proteins are categorized according to the length of the cytoplasmic domains (short or long) they have or the quantity of extracellular domains (2D or 3D) they possess. The long cytoplasmic domains trigger inhibitory signals through the immune tyrosine-based inhibition motif (ITIM) as compared with the short cytoplasmic domains which trigger activating signals through the immune tyrosine-based activation motif (ITAM) [2].

Like other aKIRs, KIR2DS2 has two functional domains namely: Ig-like C2 type 1 and Ig-like C2 type 2 domains spanning from residues 42-107 and 142-205 respectively. Within these functional domains, KIR2DS2 differs from KIR2DL2 (an inhibitory receptor) with only three residues Tyrosine 45, Arginine 148, and Threonine 200; while it differs from KIR2DL3 (an inhibitory receptor) at only one residue: Tyrosine 45. Tyrosine 45 and Glutamine 71 prevents the recognition of HLA-C ligands by KIR2DS2. KIR2DS2 transduce signals through the TYRO protein tyrosine kinase binding protein (DAP12), Zeta-chain associated protein kinase (ZAP70) and Spleen tyrosine kinase (syk) molecules [3,4].

Like other Killer cell immunoglobulin-like receptors (KIRs), KIR2DS2 regulates NK cells function and this defines their involvement in conditions such as cancers, inflammations, autoimmune disorders, and infectious diseases [1]. In autoimmunity, KIR2DS2 has been shown to influence alloreactivity and increase the number of EB6 (KIR2DL1/S1)-expressing NK clones [5]. As in other KIRs, KIR2DS2 also affects the susceptibility to various diseases such as ulcerative colitis [6], cytomegalovirus infection [7], and hepatitis C [8].

Though largely unidentified, the ligands for KIR can be found in the subsets of Human Leucocyte Antigen (HLA) class I molecules. Whereas HLA–ligand specificities are well established for iKIRs, they are not for aKIRs [9,10]. In spite of a high structural similarity, KIR2DS2 does not bind with HLA-C2 while KIR2DL2 and KIR2DL3 do [11,12]. However, a successful crystallization of KIR2DS2 in complex with HLA-A*11:01 has been reported [13]. The Tyr 45 residue of KIR2DS2 binds to the NH group of Thr80 of the α-1 helix of HLA-A*11:01 forming a hydrogen bond. This is remarkably different from iKIRs that use residue 44 to bind to Asn80 in HLA-C1 or Lys80 in HLA-C2 [13]. Beyond this, KIR2DS2 also recognizes a β2-microglobulin-independent ligand expressed on cancer cells [14].

N-acetyl-D-glucosamine (NAG) has been known to bind effectively with the activation receptors of the rat natural killer cells [15]. Similar to NAG, researchers on this project were saddled with the responsibility of prospecting for other small molecules that have an immuno-stimulatory effect on the KIR2DS2 receptor. A library of compounds from fruits and vegetables was used with the hope that the leads generated from this research would be used to produce nutraceuticals.

## 2.0 Methodology

### Materials

The ligand and protein databases used include Protein Databank, Uniprot, and PubChem. MolProbity, ExPasy, Chiron, pkCSM, Swissadme, Molinspiration, Clustal Omega, Protein-Ligand Interaction Profiler (PLIP), Galaxy and MDWEB were used as webservers. The software used were Pymol, Python prescription (PyRx) 0.8 and Discovery studio 2017.

### Methods

#### Preparation, analysis and validation of target protein structure

The crystal structure of the human natural killer cell activator receptor KIR2DS2 (PDB ID: 1m4k) was retrieved from the Protein Data Bank. The visualization tool, *Pymol* was used to remove water molecules and ligand from the protein structure [16]. The online server, *Chiron* was used for energy minimization to reduce the steric clashes of the protein structure [17] The Volume, Area, Dihedral Angle Reporter (VADAR 1.8) webserver was used to reveal the architecture of the target protein while it was analyzed and validated using the Ramanchandran and hydrophobicity plots obtained from the MolProbity and EXPasy web servers respectively [18,19].

#### Molecular docking

A library of 1,697 natural compounds was built from 79 edible African plants which are predominantly fruits, vegetables and spices. 1m4k was docked against compounds in the library that are totally complaint with Lipinski and Veber rules using the *PyRx* 0.8 version software 0.8 [20]. All ligands were converted from the SDF to pdbqt format. For stable conformation, the conjugate gradient descent was used as optimization algorithm and the Universal Force Field (UFF) was used as the energy minimization parameter.

#### Screening of Results

A reference compound, N-acetyl-D-Glucosamine was docked against the target protein and a score of −5.70 kcal/mol was obtained. All the docked complexes were further evaluated for potency using the Ligand Efficiency Metrics (LEM). The LEM used are the Ligand Efficiency (LE), ligand-efficiency-dependent lipophilicity, (LELP) and the Ligand-lipophilicity efficiency (LLE). Further screening for molar refractivity, saturation, promiscuity, pharmacokinetic properties, and bioactivity were done using the *SWISSADME, pkCSM*, and *Molinspiration* webservers respectively [21].

#### Binding Site analysis

After molecular docking, the poses of the selected ligands as they interact with the receptor were saved on *PyRx* and viewed on *PyMol*. The protein structure was superimposed on PyMol and saved in the pdb format. The resultant structure was uploaded into the Protein-Ligand Interaction Profiler (PLIP) webserver to reveal the three-dimensional depiction of best-docked complex as shown by the binding site and all the protein-ligand interactions [22,16]. Compounds that had no hydrogen bond interaction with TYR45 were eliminated. However, a front runner compound, Monocrotalline which had hydrophobic interaction at TYR 45 was eliminated because of its established toxicity profile. Though computationally predicted to be safe, Monocrotaline is used to induce Pulmonary arterial hypertension in experimental animals [23]

#### Molecular Dynamics Simulations

The molecular dynamics simulation (MDS) of the native protein and all its ligand complexes using the Galaxy (versions 2019.1 and 2019.1.4) supercomputing server which uses the Groningen Machine for Chemical Simulations (GROMACS) software [24] GROMACS-compatible Gro and topology files for small molecules were generated using *LigParGen* with OPLS-AA/1.14*CM1A force field parameters [25]

The parameters for the initial simulation set up include: SPC/E for water model; OPLS/AA as the force field; box type is rectangular with all equal sides each measuring 1.0nm; and Hydrogen was ignored. GROMACS solvation and addition of ions was done using the SPC water model. Energy minimization was conducted using the steepest descent algorithm for 50,000 steps. 1.0 was used for Distance cut-offs include for short range van der Waals, Coulomb, and the shortrange neighbor list respectively. Other parameters for minimization include Fast smooth Particle-Mesh Ewald electrostatics and EM tolerance of 1000.Equilibration was performed using NVT and NPT ensembles with the following parameters: leap-frog integrator, constrained Bonds with H-atoms, 300k temperature, 0.002 ps step length and 5,000 steps between saving data points. Finally, a 1ns molecular dynamics simulation was carried out for the Apo and Holo proteins with 500,000 steps.

Using the BIO3D tool on the Galaxy super-computing platform, the Root Mean Square Deviation of atomic positions (RMSD), Root Mean Square Fluctuation (RMSF) of protein backbone, Dynamical Cross-Correlation Matrix (DCCM), Principal Component Analysis (PCA), were determined [26]. The Radius of Gyration (RoG) and B Factor were obtained using the MDWEB webserver [27].

## 3.0 Discussion of Results

### Structural analysis, validation and preparation of KIR2DS2 (PDB ID: 1m4k)

Figure 1 reveals the Kyte & Doolittle hydrophobicity plot of the full protein structure as found in the UniProt database (P43631). The structure consists of signal peptides (1-21), extracellular domains (22-245), helical domain (246-265), and cytoplasmic domain (266-304). Within the extracellular domain are the Ig-like C2-type 1 (42-107) and Ig-like C2-type 2 (142-205) subdomains. From Figure 2, the signal peptide is hydrophilic while Ig-like C2-type 1 and Ig-like C2-type 2 domains are predominantly hydrophobic.

**Figure 1:**
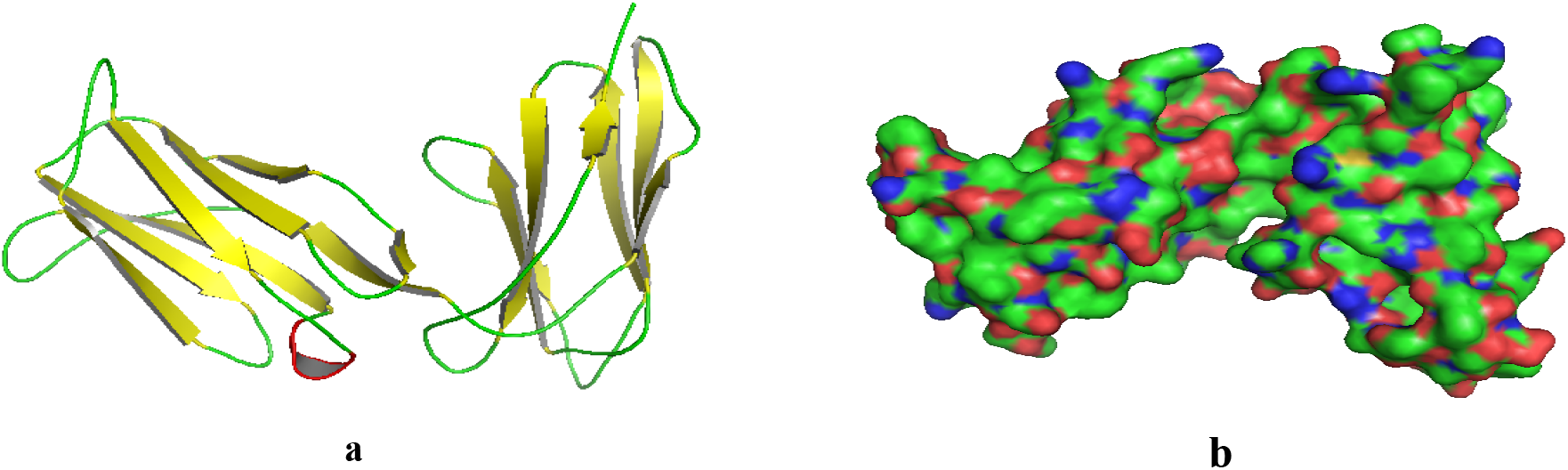
a: Cartoon model of the **c**rystal structure of KIR2DS2 (PDBID: 1m4k): Beta sheets (yellow), Alpha helix (red) and Loops (green) b: Surface representations.

**Figure 2:**
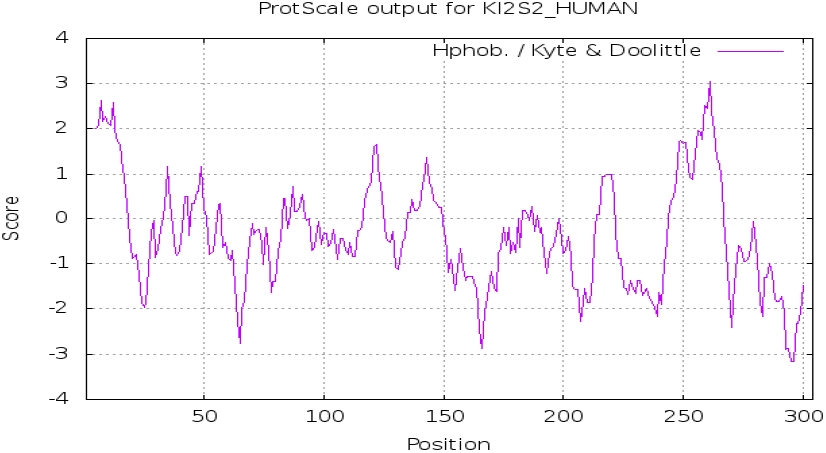
Kyte & Doolittle hydrophobicity plot of KIR2DS2 (PDBID: 1m4k)

The Apo structure, 1m4k is the extracellular domain of KIR2DS2 (Figure 1). It has 193 amino acids with the following constituent secondary structures: α helix 0 %; beta sheets 72%; Coil 27%; and Turns 10%. Total Accessible Solvent Area (ASA) is 9074.7(Å)^2^ X ray diffraction study revealed resolution is 2.3Å and unit cell crystal dimensions are a = 97.535 Å, b = 97.535 Å, and c = 54.375 Å for α (90°), β (90°) and γ (120°) angles respectively. The R-Value is 0.221 and Rfree value is 0.247.

From Figure 3, Ramachandran analysis of the candidate structure shows that 95.29% of residues are within the favoured region and 2.09% outliers. Rotamer analysis shows 91.93% of rotamers are within the favoured region and 4.97% poor rotamers. Further protein geometry reveals no Cβ deviations, 0.13% bad bonds and 0.24% bad angles. The VDW repulsion energy of 1m4k was minimized by *Chiron* from 56.15 Kcal/mol (clash ratio of 0.023) to 35.85 Kcal/mol (clash ratio of 0.015).

**Figure 3:**
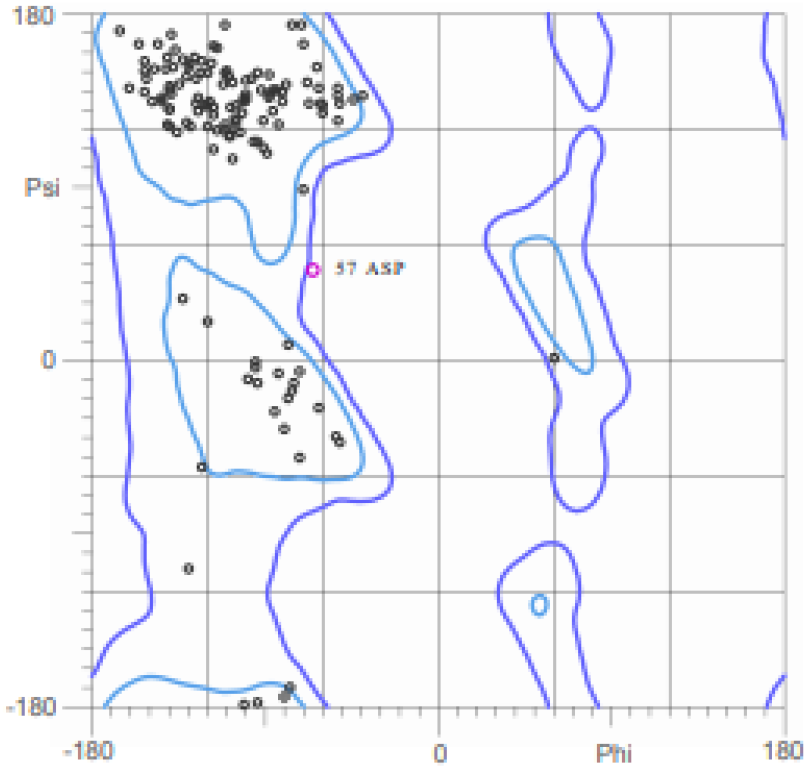
Ramanchandran plot for KIR2DS2 (PDBID: 1m4k)

### Chemoinformatic profile of ligands

(Figure 4, Table 1): Drug-like properties are important for receptor-binding, bioavailability and cellular uptake within the body. To meet these criteria, all the molecular descriptors of the frontrunner compounds should not violate the Lipinski (RO5), Veber, and Ghose rules. Put together these rules state that the molecular weight should be ≤ 500g/mol; hydrogen bond acceptors should be ≤ 10; hydrogen bond donors should be ≤ 5; Log P should be ≤ 5; the polar surface area should be ≤ 140A^2^; number of rotatable bonds should be < 10; and molar refractivity should be between 40-130 cm3 [28,29,30].

**Figure 4:**
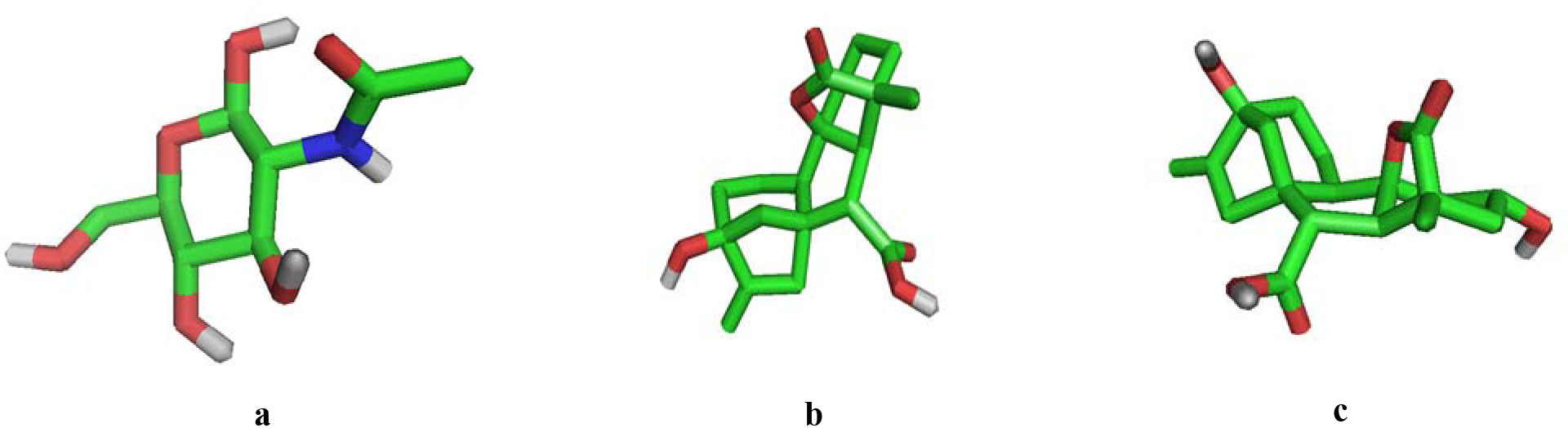
The 3D chemical structures (stick model) of standard and lead compound a: NAG b: GA20 c: GA29

**Table 1:** Chemo-informatic properties of standard and lead compound

|  | **NAG (standard)** | **GA20** | **GA29** |
| --- | --- | --- | --- |
| Molecular Formula | C8H15NO6 | C19H24O5 | C19H24O6 |
| Molecular Weight<br>(g/mol) | 221.21 | 332.4 | 348.4 |
| Log P | -1.7 | 1.2 | 0.2 |
| Hydrogen Bond<br>Acceptors | 5 | 5 | 6 |
| Hydrogen Bond Donors | 5 | 2 | 3 |
| # heavy atoms | 15 | 24 | 25 |
| # rotatable bonds | 2 | 1 | 1 |
| TPSA (Ål) | 119 | 83.8 | 104 |
| Molar Refractivity | 47.19 | 86.18 | 87.34 |
| Saturation (fraction csp<sup>3</sup>) | 0.88 | 0.79 | 0.79 |
| PAIN Alert | No | No | No |
| GCPR ligand | -0.47 | 0.22 | 0.24 |
| Ion channel modulator | -0.30 | 0.23 | 0.20 |
| Kinase Inhibitor | -0.71 | -0.21 | -0.24 |
| Nuclear receptor ligand | -0.74 | 0.49 | 0.60 |
| Protease inhibitor | 0.07 | 0.09 | 0.19 |
| Enzyme inhibitor | 0.46 | 0.30 | 0.42 |

Our results prove that GA20, GA29 and the standard did not violate the Lipinski (RO5), Veber, and Ghose rules indicating potentially good drug permeability. Specifically, the standard has the highest lipophilicity of all the compounds suggesting that it has the greatest absorbability across lipid membranes. While the standard and GA 29 will not permeate the blood-brain barrier, GA 20 will because its TPSA value is less than 90 angstroms squared [31].

In computational drug discovery, the molecular complexity of organic molecules can be measured by saturation. GA20, GA29 and the standard have their fraction of carbons in the sp3 hybridization more than 0.25(Table 1). The standard has the highest saturation [32]. PAINS (pan assay interference) or promiscuous compounds are frequent hitters that contain potentially problematic moieties that yield false positive response in biological assays. GA20, GA29 and the standard had no PAIN alerts [33].

Further analysis revealed that all the compounds showed good lead-like behavior with good predicted bioactivity scores against the major drug targets such as GCPRs, ion channels, kinases, nuclear receptors, proteases and enzymes. GA20, GA29 and the standard show moderate kinase inhibition activity. Predictably for all the targets except enzyme inhibition, GA20 and GA29 showed greater activity than the standard. Of the three compounds, the standard has the best enzyme inhibition value [34]. Overall, the results show that GA20, GA29 and the standard are good drug candidates (Table 1).

### Pharmacokinetic properties of ligands

Novel drugs should have good pharmacokinetic properties. The Absorption, Distribution, Metabolism, Excretion, and Toxicity (ADMET) properties of the front-runner compounds were predicted *in silico* using graph-based signatures (Table 2).

**Table 2:** Pharmacokinetic properties of standard and lead compounds

|  | **NAG** | **GA20** | **GA29** |
| --- | --- | --- | --- |
| Water solubility (log mol/L) | -1.38 | -2.636 | -2.659 |
| Caco2 permeability (log Papp in 10<sup>-6</sup> cm/s) | -0.219 | 1.186 | 0.691 |
| Human Intestinal absorption (% Absorbed) | 31.963 | 98.911 | 71.42 |
| Skin Permeability (log Kp) | -3.234 | -2.735 | -2.735 |
| P-glycoprotein substrate (Yes/No) | No | No | No |
| P-glycoprotein I inhibitor (Yes/No) | No | No | No |
| P-glycoprotein II inhibitor (Yes/No) | No | No | No |
| VDss (human) (log L/kg) | 0.041 | -0.829 | -0.82 |
| Fraction unbound (human) (Fu) | 0.856 | 0.418 | 0.466 |
| BBB permeability (log BB) | -0.618 | -0.209 | -0.601 |
| CNS permeability (log PS) | -3.694 | -2.998 | -3.112 |
| CYP2D6 substrate (Yes/No) | No | No | No |
| CYP3A4 substrate (Yes/No) | No | Yes | Yes |
| CYP1A2 inhibitor (Yes/No) | No | No | No |
| CYP2C19 inhibitor | No | No | No |
| (Yes/No) |  |  |  |
| CYP2C9 inhibitor<br>(Yes/No) | No | No | No |
| CYP2D6 inhibitor<br>(Yes/No) | No | No | No |
| CYP3A4 inhibitor<br>(Yes/No) | No | No | No |
| Total Clearance (log<br>ml/min/kg) | 0.711 | 0.417 | 0.42 |
| Renal OCT2 substrate<br>(Yes/No) | No | No | No |
| AMES toxicity (Yes/No) | No | No | No |
| Max. Tolerated dose<br>(human) (log mg/kg/day) | 1.944 | 0.371 | 0.262 |
| hERG I inhibitor<br>(Yes/No) | No | No | No |
| hERG II inhibitor<br>(Yes/No) | No | No | No |
| Oral Rat Acute Toxicity<br>(LD50) (mol/kg) | 1.547 | 2.051 | 2.097 |
| Oral Rat Chronic Toxicity<br>(log mg/kg_bw/day) | 3.406 | 2.135 | 2.498 |
| Hepatotoxicity<br>(Yes/No) | No | No | No |
| Skin Sensitization<br>(Yes/No) | No | No | No |
| *T.Pyriformis* toxicity (log<br>ug/L) | 0.285 | 0.285 | 0.285 |
| Minnow toxicity (log<br>mM) | 4.705 | 1.958 | 2.694 |

Absorption parameters such as water solubility (insoluble: −4.0 Log mol/L), caco2 permeability (high:> 0.9), human intestinal absorption (poor: <30%), and skin permeability (low: LogKp >-2.5) are indicators of the therapeutic potential of chemical compounds as these ensure their penetration to reach the target molecule. All values obtained for GA20, GA29 and the standard are within pharmacological range and none of them are inhibitors of p-glycoprotein [35,36].

Comparing to standard values, distribution properties such as fraction unbound (not less than 0.1), volume of distribution (Low: Log VDss <- 0.15; High: Log VDss > 0.45), the Blood Brain Barrier (permeable: Log BBB > 0.3; poor <: Log BBB <-1), and Central Nervous System (permeable Log PS > −2; poor Log PS < −3) permeability were evaluated. All values obtained for GA20, GA29 and the standard for volume of distribution and the Blood Brain Barrier are within pharmacological range. However, the standard has a poor CNS permeability [35,37].

Inhibition of some isoforms of cytochrome P450 could make a novel molecule toxic. The predicted metabolic behavior of GA20, GA29 and the standard shows no inhibition of CYP1A2, CYP2C19, CYP2C9, CYP2D6 and CYP3A4 enzymes. However, GA20 and GA29 are substrates of CYP3A4. This means that their dose would be affected either by induction or the inhibition of CYP3A4 [38].

The predicted excretion values for Total Clearance for GA20, GA29 and the standard are within pharmacological range [39]. GA20, GA29 and the standard are not substrates of Renal Organic Cation Transporter 2 (OCT2). This implies that Renal OCT2 will eliminate none of them from the blood into the proximal tubular cell [40].

The maximum recommended tolerated dose determines the dose to be administered in the phase 1 of clinical trials. GA20 and GA29 have low maximum recommended tolerated dose (less than 0.477 log mg/kg/day) while the standard has a high value (more than 0.477 log mg/kg/day). GA29 is the most potent [35].

The toxicity required to inhibit 50% of the growth of *T.pyriformis* (IGC50), protozoan bacteria is very important parameter measured in drug discovery. A compound is considered toxic when the pIGC50 (negative logarithm of the concentration required to inhibit 50% growth in logUg/L) value is greater than −0.5 log Ug/L. From the results, GA20, GA29 and the standard are predicted to be toxic against *T.pyriformis*. This suggests an antibacterial effect and might not be toxic to human cells [35]. Similarly, log LC50 is the log of a compound to cause death of 50% of flathead Minnows. A log LC50 value less than 0.3 log mM signifies high acute toxicity. From the results, GA20, GA29 and the standard are not toxic to Minnows [35].

HerG inhibition and AMES toxicity are critical toxicity parameters as it reveals the cardio-toxic properties and mutagenic behavior of the compounds respectively. GA20, GA29 and the standard did not show HerG inhibition, or AMES toxicity. They all also did not show dermatotoxic and hepatotoxic properties [35].

### Molecular docking analyses of ligands against KIR2DS2

Ligand Efficiency Metrics (LEM) was used to analyze potency the compounds based on their size and lipophilicity [41]. The LEM used include Lipophilic Efficiency (LE), Ligand-Efficiency dependent Lipophilicity (LEDL), and Ligand Lipophilicity Efficiency (LLE). As seen in Table 3, GA20, GA29 and the standard met the cut-off LE (≥ 0.3), LELP (−10 to 10) and LLE (greater than 5) [42]. The standard and the lead compounds met the cut-offs for all the LEM showing the required potency.

**Table 3:** Molecular docking scores and Ligand Efficiency Metrics of ligands against KIR2DS2

| **Ligand** | Binding (Kcal/mol)<br>affinity | **LE** | **LELP** | **LLE** |
| --- | --- | --- | --- | --- |
| N-acetyl-D-Glucosamine | -5.70 | 0.38 | -4.47 | 7.4 |
| GA20 | -7.40 | 0.31 | -3.87 | 6.2 |
| GA29 | -8.20 | 0.33 | -0.61 | 8.0 |

### Binding Site analyses

Hydrogen bond determines the specificity of ligand binding and it captures two key factors which are the length and orientation of the bond [43]. From Figures 5 & 6 and Table 4, the standard and lead compounds all have hydrogen bonding with residues within the functional domains of the target which are the Ig-like c2 type 1 (42-107) and the Ig-like c2 type 2 (142-205) domains. NAG had the highest number of hydrogen bonds within these domains.

**Figure 5:**
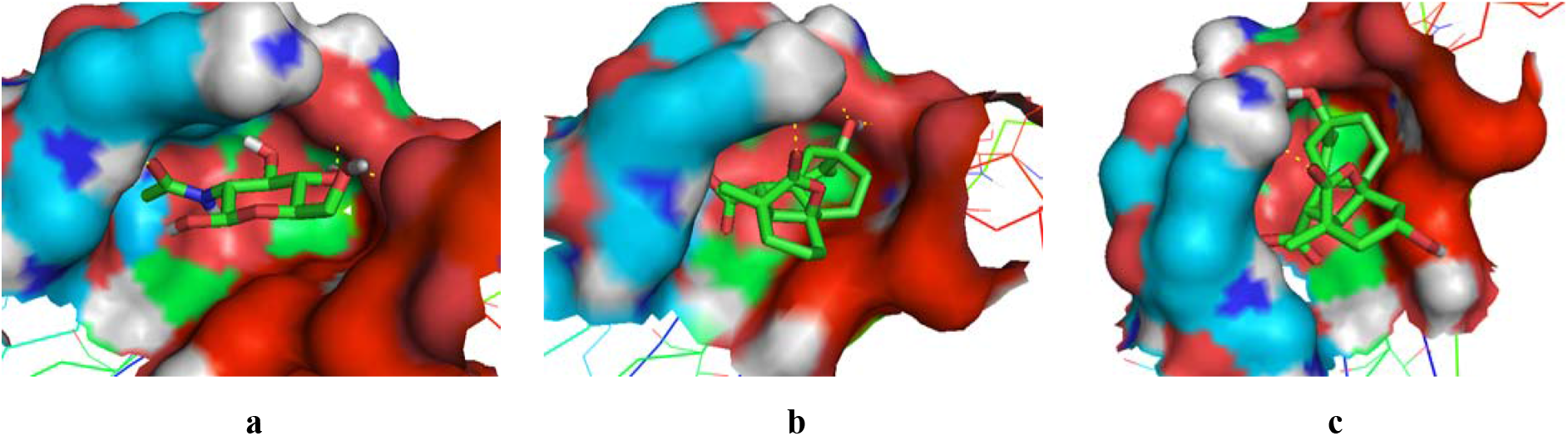
Binding site of KIR2DS2 interacting with standard and lead compounds a: KIR2DS2 – NAG b: KIR2DS2-GA20 c: KIR2DS2-GA29

**Figure 6:**
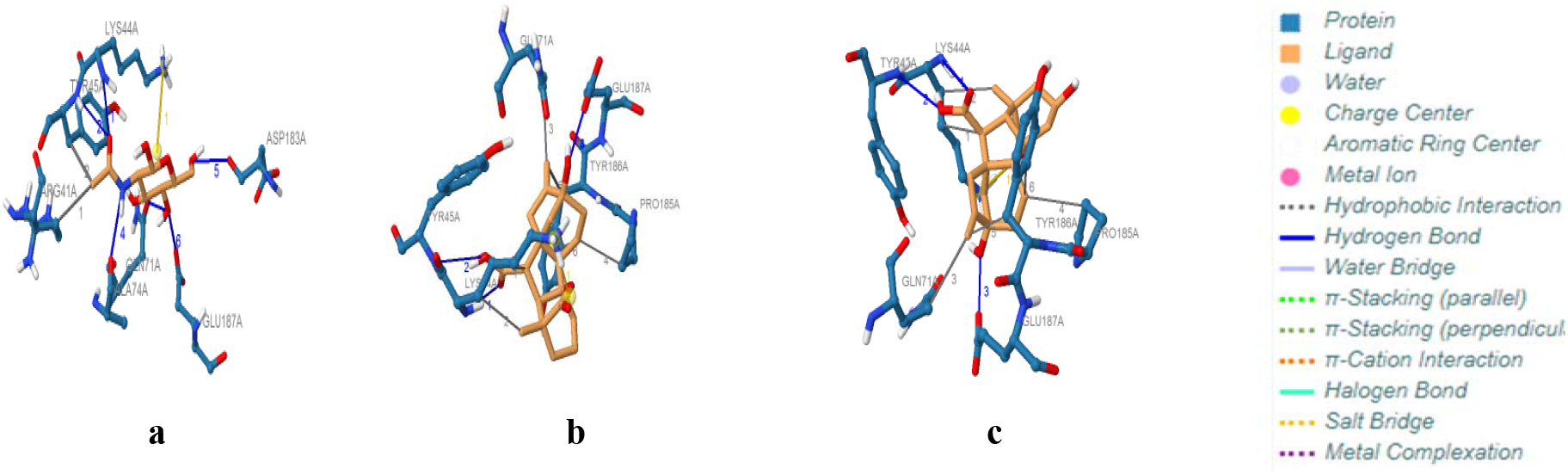
Protein-Ligand interactions of KIR2DS2 with standard and lead compound a: KIR2DS2-NAG complex b: KIR2DS2-GA20 complex c: KIR2DS2-GA29 complex

**Table 4:** Hydrogen bond analysis of KIR2DS2 with standard and lead compounds

| **Compound** | **Number of bonds** | **Residues** | **Distance (H-A)** | **Distance (D-A)** | **Bond angle** |
| --- | --- | --- | --- | --- | --- |
| N-acetyl-D-Glucosamine (Standard) | 6 | LYS 44 | 2.32 | 2.88 | 113.53 |
|  |  | TYR45 | 2.65 | 3.15 | 110.26 |
|  |  | GLN 71 | 2.37 | 2.94 | 115.97 |
|  |  | ALA74 | 2.15 | 3.14 | 161.14 |
|  |  | ASP183 | 2.29 | 3.09 | 136.96 |
|  |  | GLU187 | 2.01 | 2.70 | 129.79 |
| GA 20 | 4 | LYS 44 | 3.14 | 3.93 | 135.58 |
|  |  | TYR45 | 2.88 | 3.40 | 112.2 |
|  |  | GLU187 | 2.69 | 3.10 | 106.90 |
|  |  | GLU187 | 2.19 | 3.10 | 151.21 |
| GA29 | 3 | LYS44 | 3.15 | 3.95 | 136.17 |
|  |  | TYR45 | 2.90 | 3.42 | 111.94 |
|  |  | GLU187 | 2.71 | 3.10 | 105.81 |

Notably, the standard and the lead compounds form hydrogen bonds on LYS44 and TYR45. Only NAG forms hydrogen bond at GLN 71. These residues are essential in that they affect the binding of the KIR2DS2 differs from KIR2DL2 (an inhibitory receptor) with HLA molecules. This suggests that as these compounds can both trigger the two receptors [3,4,13]. In terms of the donor to acceptor distance, NAG forms a moderate (2.5-3.2 Å) hydrogen bond at TYR 45 while GA 20 and GA29 form weak (3.2-4.0 Å) hydrogen bonds at the same residue. In spite of the different strata of hydrogen bond strength, only a slight difference exists between the standard (3.15 Å) and GA 20 and GA29 (3.40 and 3.42Å respectively). In terms of bond angle formed at TYR45, all compounds (standard and the leads) form weak (less than 130°) hydrogen bonds. [44,45].

From Figure 5 and Table 5, all the compounds form other kinds of interactions at LYS44. The standard forms a salt bridge at this residue while the lead compounds form both salt bridges and hydrophobic interactions. The presence of salt bridges in all the protein-ligand interactions of the standard and lead compounds enhances the strength and stability of the complexes [46]. The strength of the salt bridge is affected by the distances between the atoms of ligand and those of residues. The distance required is less than 4 Å [47]. This suggests that the salt bridge formed by NAG at LYS44 is a weak one.

**Table 5:**
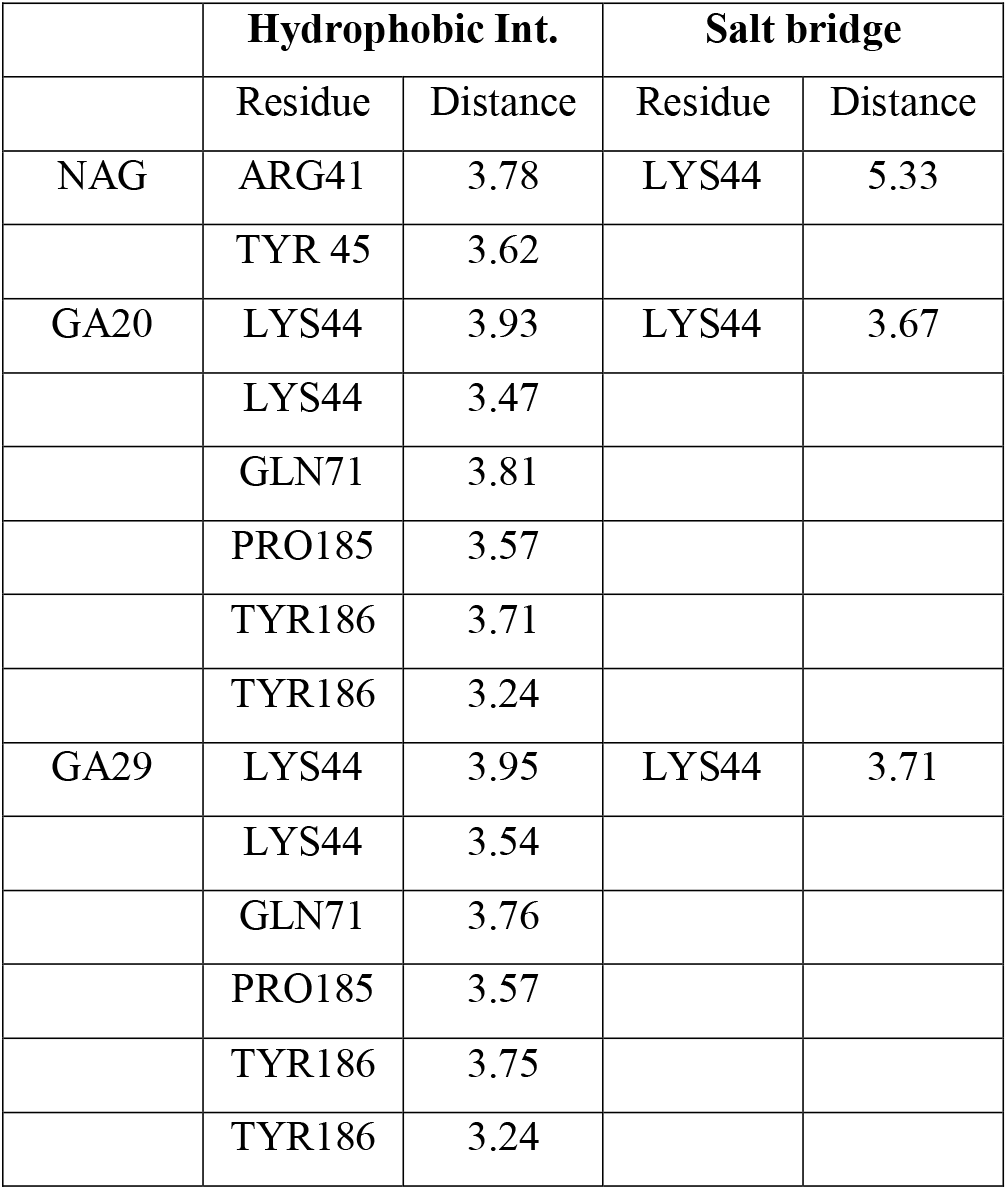
Other Protein-ligand interactions.

|  | Hydrophobic Int. |  | Salt bridge |  |
| --- | --- | --- | --- | --- |
|  | Residue | Distance | Residue | Distance |
| NAG | ARG41 | 3.78 | LYS44 | 5.33 |
|  | TYR 45 | 3.62 |  |  |
| GA20 | LYS44 | 3.93 | LYS44 | 3.67 |
|  | LYS44 | 3.47 |  |  |
|  | GLN71 | 3.81 |  |  |
|  | PRO185 | 3.57 |  |  |
|  | TYR186 | 3.71 |  |  |
|  | TYR186 | 3.24 |  |  |
| GA29 | LYS44 | 3.95 | LYS44 | 3.71 |
|  | LYS44 | 3.54 |  |  |
|  | GLN71 | 3.76 |  |  |
|  | PRO185 | 3.57 |  |  |
|  | TYR186 | 3.75 |  |  |
|  | TYR186 | 3.24 |  |  |

The number of hydrophobic interactions is three times higher in GA20 and GA29 as compared with the standard. These optimal interactions suggest that GA 20 and GA29 are have more atom efficient binding than NAG [48]. Only NAG had a hydrophobic interaction at TYR 45, while the lead compounds had at GLN 71. While TYR45A is important in determining the activating signal of the target; GLN 71, is in determining whether the target binds to HLA-C [13,3,4].

## Molecular Dynamic Simulation Analyses

### Structures

Comparing the crystal structure with the simulated apo and holo structures suggests that there is an unfolding of the alpha helix at residues 64, 65 and 66 during the molecular dynamic simulation (Figure 1 and 7).

**Figure 7:**
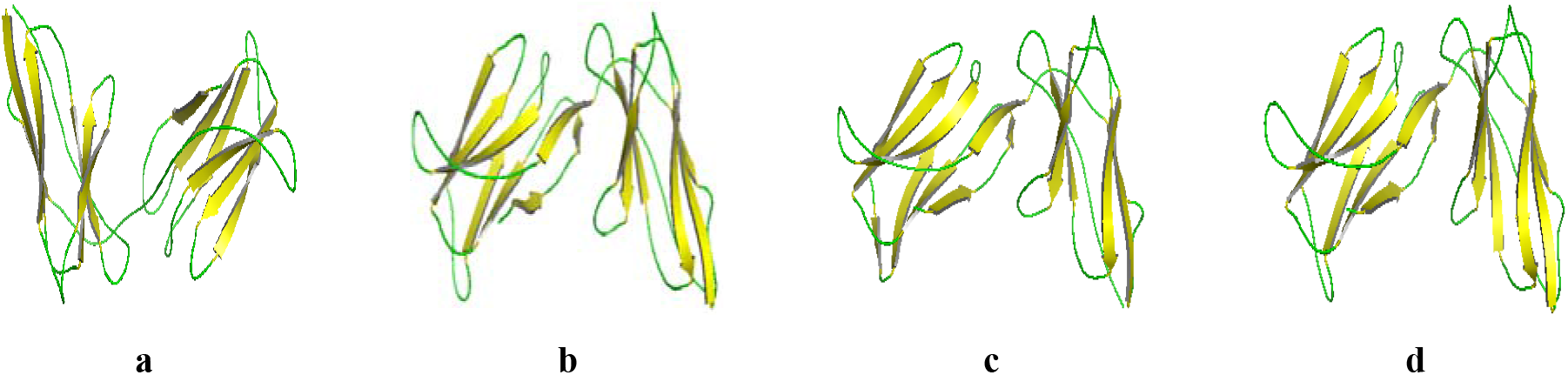
Cartoon model of the crystal structure of KIR2DS2 Apo and Holo structures (without water and ions) after molecular dynamics simulation. Beta sheets (yellow), Alpha helix (red) and Loops (green) a: KIR2DS2; b: KIR2DS2-NAG c: KIR2DS2-GA20 d: KIR2DS2-GA29

### Root Mean Square Deviation of Atomic Positions (RMSD)

**RMSD** is the most common quantitative measure of similarity between two superimposed protein structures (the reference and target structures). It measures the variations in the distances between atoms in two superimposed structures and an RMSD value of 0.0 indicates a perfect overlap [49].

From Figure 8 and Table 6, the RMSD of the simulated Apo protein relative to the crystal structure suggests a gradual increase as the production time increased. It peaked at Frame 43 (2.51 Å) and thereafter stabilized. The global RMSD for the Apo structure is 180.94 Å while the average is 1.79 Å. The RMSD for the simulated KIR2DS2 – NAG complex suggests a steady rise until it peaked at Frame 70 (2.25 Å) and thereafter stabilized. The global RMSD for this holo structure is 174.91 Å while the average is 1.73 Å. The RMSD for the simulated KIR2DS2-GA20 complex suggests a steady rise until it peaked at Frame 39 (2.26 Å) and thereafter stabilized. The global RMSD for this holo structure is 166.41 Å while the average is 1.65 Å. Also, the RMSD for the simulated KIR2DS2 – GA29 complex suggests a steady rise until it peaked at Frame 65 (2.5 Å) and thereafter stabilized. The global RMSD for this holo structure is 165.22 Å while the average is 1.64 Å. From Figure 9 and Table 6, the RMSD histogram suggests that peak distribution is more tilted to the left in the KIR2DS2 – GA20 and KIR2DS2 – GA29 complexes than the KIR2DS2 – NAG complex. Specifically, KIR2DS2 – GA29 had the highest number of peaks within the 1.0 to 1.5 Å range.

**Figure 8:**
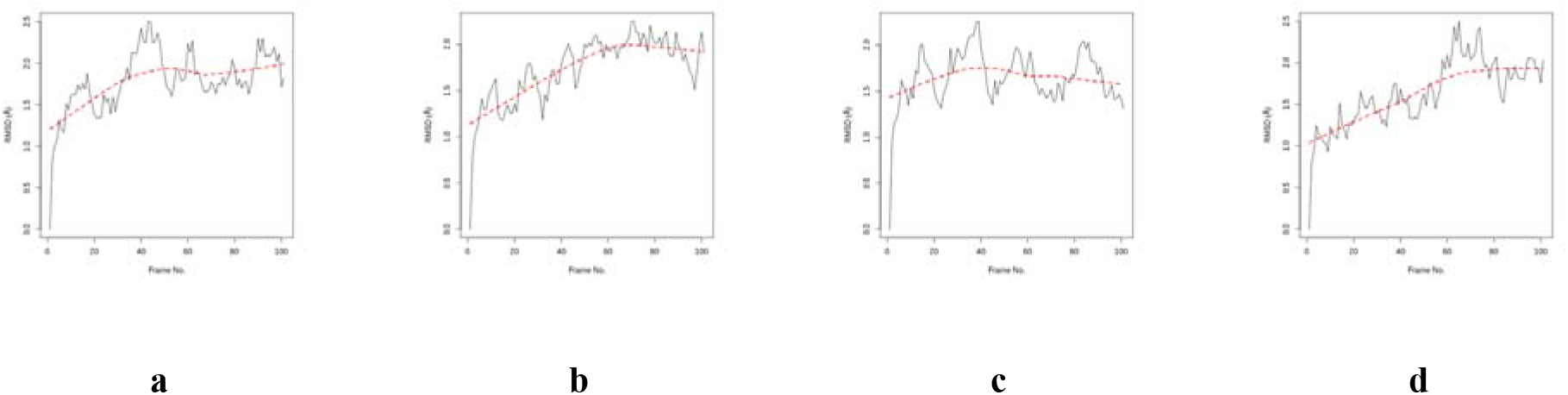
RMSD for Apo and Holo structures (a). KIR2DS2 (b) KIR2DS2-NAG (c) KIR2DS2-GA20 (d) KIR2DS2-GA29

**Fig 9:**
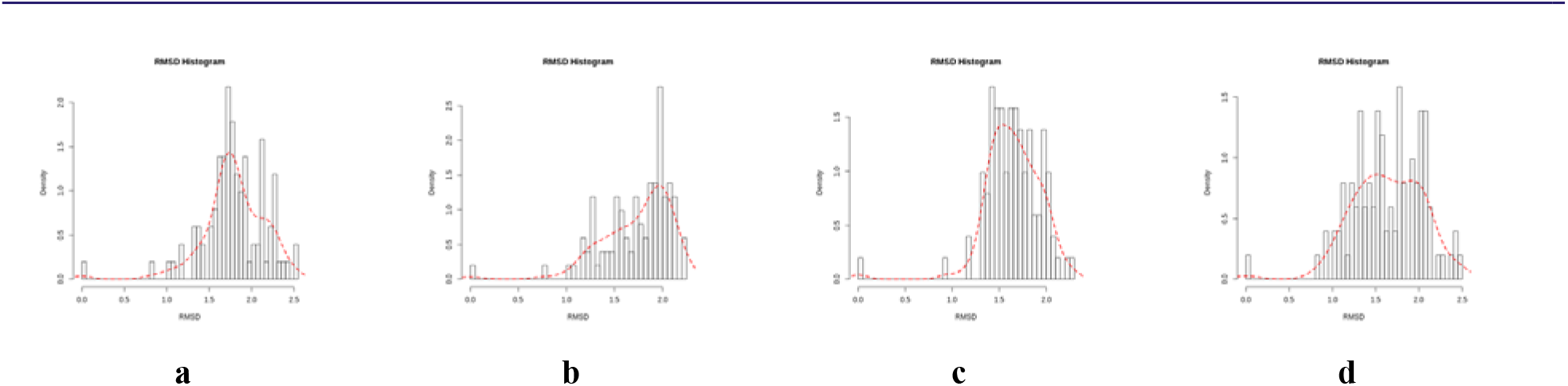
RMSD histogram for Apo and Holo structures (a). KIR2DS2 (b) KIR2DS2-NAG (c) KIR2DS2-GA20 (d) KIR2DS2-GA29

**Table 6:** Summary of data from Molecular Dynamics Simulations of Apo and Holo structures.

| **MDS Parameters** | **KIR2DS2** | **KIR2DS2 + NAG** | **KIR2DS2 + GA20** | **KIR2DS2 + GA29** |
| --- | --- | --- | --- | --- |
| **RMSD** |  |  |  |  |
| Total RMSD | 180.94 | 174.91 | 166.41 | 165.22 |
| Average RMSD | 1.79 | 1.73 | 1.65 | 1.64 |
| Lowest RMSD | 0.00 | 0.00 | 0.00 | 0.00 |
| Highest RMSD | 2.51 | 2.25 | 2.26 | 2.50 |
| Time Frame of Highest RMSD | 43 | 70 | 39 | 65 |
| Time Frame of Lowest RMSD | 1 | 1 | 1 | 1 |
| **RMSD Peak Distribution** |  |  |  |  |
| 0.00 - 0.50A | 1 | 1 | 1 | 1 |
| 0.50 - 1.00A | 1 | 1 | 1 | 3 |
| 1.00 - 1.50A | 12 | 20 | 27 | 33 |
| 1.50 - 2.00A | 60 | 55 | 62 | 41 |
| 2.00 - 2.50A | 25 | 24 | 10 | 22 |
| 2.50 - 3.00A | 2 | 0 | 0 | 1 |
| **RMSF** |  |  |  |  |
| Total Global RMSF | 165.32 | 170.97 | 147.41 | 157.00 |
| Average Global RMSF | 0.86 | 0.89 | 0.76 | 0.81 |
| Total Regional (Res 42-107) RMSF | 57.35 | 58.35 | 50.30 | 49.52 |
| Average Regional (Res 42-107) RMSF | 0.87 | 0.88 | 0.76 | 0.75 |
| Local RMSF (Res. 45) | 0.78 | 0.94 | 0.78 | 0.75 |
| Least Fluctuation | 0.40 | 0.41 | 0.37 | 0.36 |
| Highest Fluctuation | 2.20 | 2.26 | 1.46 | 2.15 |
| Range of RMSF | 1.80 | 1.85 | 1.09 | 1.79 |
| **Radius of Gyration** |  |  |  |  |
| Average Gyration | 4.0377 | 4.0377 | 4.0349 | 4.0329 |
| Minimum Gyration | 4.0270 | 4.0248 | 4.0264 | 4.02575 |
| Maximum Gyration | 4.0469 | 4.0426 | 4.0431 | 4.04089 |
| Range of Gyration | 0.0199 | 0.0178 | 0.0167 | 0.01514 |
| % Gyration | 0.49 | 0.44 | 0.41 | 0.38 |
| Time Frame of Maximum Gyration | 65 | 55 | 60 | 74 |
| Time Frame of Minimum Gyration | 1 | 1 | 15 | 13 |
| **B-Factor** |  |  |  |  |
| Global Average B Factor | 170.12 | 236.89 | 150.26 | 130.13 |
| Regional (Res. 42-107) Average B Factor | 166.78 | 248.35 | 150.38 | 123.78 |
| Local B factor (residue 45) | 191.44 | 315.63 | 138.38 | 154.77 |
| **PCA** |  |  |  |  |
| Total motions (PC1) | 11.70 | 12.01 | 12.48 | 11.43 |
| Average motions (PC1) | 0.06 | 0.06 | 0.06 | 0.06 |
| Best global Conformation | PC1 | PC2 | PC3 | PC3 |
| Best local Conformation (Residue 45) | PC1 & PC3 | PC3 | PC3 | PC3 |
| Av. Motion at Residue 45 | 0.06 | 0.06 | 0.07 | 0.05 |
| PC1 Eigenvalue | 36.30% | 49.16% | 20.94% | 36.97% |
| PC2 Eigenvalue | 12.00% | 11.26% | 17.16% | 19.94% |
| PC3 Eigenvalue | 8.90% | 6.80% | 10.04% | 6.94% |
| Total | 57.21% | 67.22% | 48.14% | 63.85% |
| PC1 cosine content | 0.91 | 0.07 | 0.85 | 0.53 |
| PC2 cosine content | 0.84 | 0.06 | 0.69 | 0.31 |
| PC3 cosine content | 0.47 | 0.12 | 0.03 | 0.29 |

Put together, 98.7% all the RMS distances between alpha carbon atoms of the three holo structures fall within the resolution of the crystal structure of the KIR2DS2 (2.3 Å) [50]. KIR2DS2 – GA20 and KIR2DS2 – GA29 complexes showed better stability than the than the KIR2DS2 – NAG complex. The KIR2DS2 – GA29 complex showed the best stability as seen with the least total and average RMSD and the highest number of peaks within the 1.0 to 1.5 Å range (greatest left shift).

### Root Mean Square Fluctuations (RMSF)

At a wide range of time scales, proteins undergo structural fluctuations under physiological and pathological conditions. The RMSF reveals the dynamic behaviour of various residues by measuring the lowest energy modes of fluctuations around their equilibrium conformations [51].

From Figure 10 and Table 6, the total and average global RMSF is smaller in the KIR2DS2 – GA20 and the KIR2DS2 – GA29 complexes than the KIR2DS2 – NAG complex showing greater stability. The KIR2DS2 – GA20 complex shows slightly lesser fluctuations globally than the KIR2DS2 – GA29 complex. In a similar manner, the total and average fluctuations within the Ig-like c2 type 1 functional domain (residues 42-107), is smaller in the KIR2DS2 – GA20 and the KIR2DS2 – GA29 complexes than the KIR2DS2 – NAG complex showing that the lead compounds induce greater stability than the standard. In this domain. The KIR2DS2 – GA29 complex shows a marginally lesser fluctuation than the KIR2DS2 – GA20 complex. Furthermore, on TYR45, both lead compounds induced less fluctuation as compared with the standard. The KIR2DS2 – GA29 complex had the least fluctuation value at this important residue.

**Figure 10:**
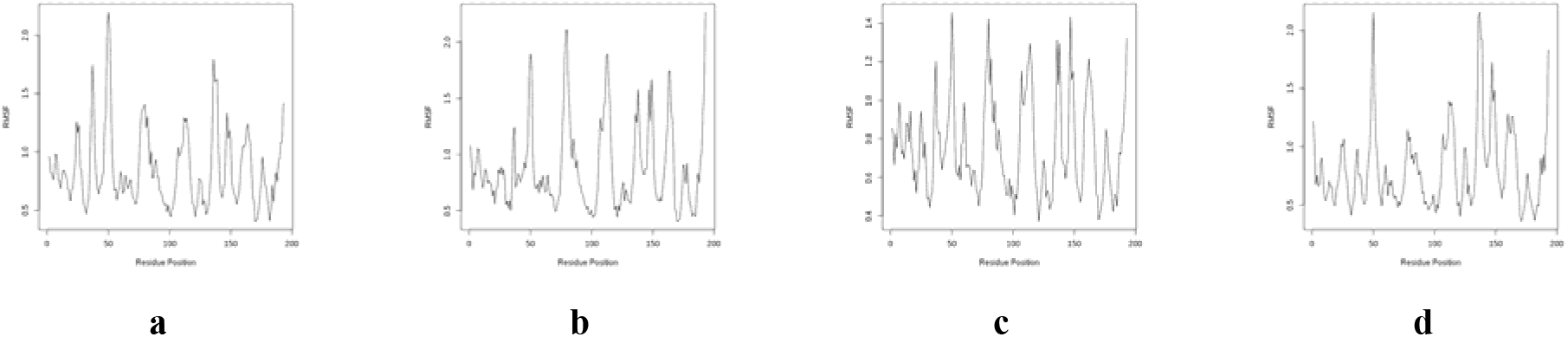
Per-residue RMSF for Apo and Holo structures (a). KIR2DS2 (b) KIR2DS2-NAG (c) KIR2DS2-GA20 (d) KIR2DS2-GA29

Overall, while evolutionary conservation of backbone fluctuations is seen in KIR2DS2, greater number of conformations due to ligand binding reveals that KIR2DS2 + GA29 complex is the most stable holo structure. Though KIR2DS2 – GA20 complex had the least global fluctuations, KIR2DS2-GA29 complex had the least regional and local (TYR 45) fluctuations.

### Radius of gyration (RoG)

The RoG is an indicator of the compactness of the secondary structures within the 3D structure of the protein. Measured from the centre of mass of the molecule, a high RoG suggests loose packing while a low RoG suggests a tight packing of the protein. [52]. After an MDS of 1ns, both Apo and holo structures showed relative compactness in terms of the degree of folding due to the hydrophobic effect and the architecture of target protein is largely conserved. This means that ligand induced conformation by NAG, GA20 and GA29 had little effect in unfolding the target protein. This is revealed in the marginal differences in their percentages of gyration. The KIR2DS2-GA29 is the most compact of all the structures. This is as suggested by the lowest average gyration over the trajectory, the lowest range of gyration and the lowest percentage gyration. Predictably, both lead compounds have less impact on the unfolding of KIR2DS2 than the standard (Figure 11 and Table 6).

**Figure 11:**
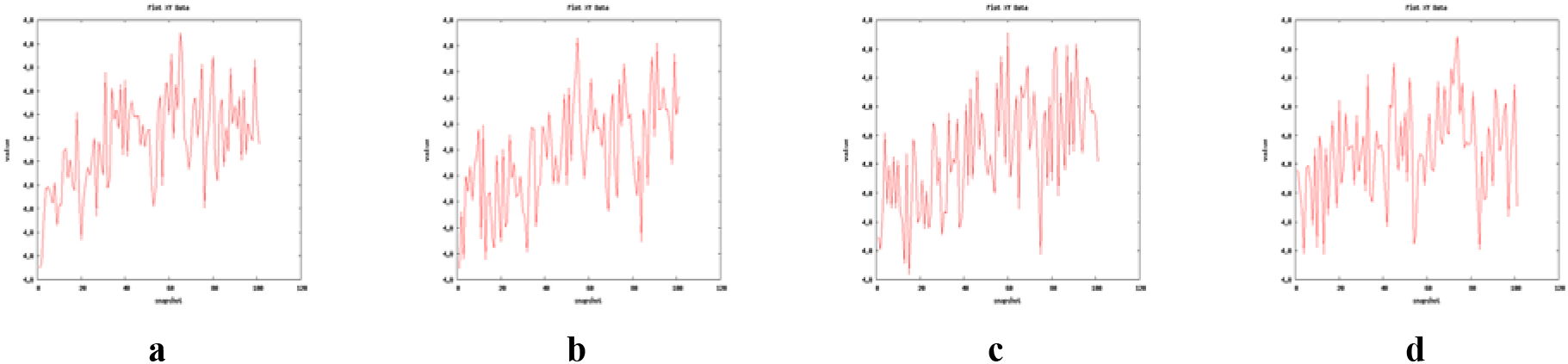
Radius of gyration for Apo and Holo structures (a). KIR2DS2 (b) KIR2DS2-NAG (c) KIR2DS2-GA20 (d) KIR2DS2-GA29

### B-Factor

This describes the thermostability of a protein molecule by identifying regions of flexibility, rigidity and internal motions [53]. From Figure 12 and Table 6, the values of the global average, regional (residues 42-107) average and local B-factors are less in the KIR2DS2-GA20 and KIR2DS2-GA29 complexes than in the KIR2DS2-NAG complex. This suggests that the lead compounds-induced conformations are more stable than the conformation induced by the standard. While KIR2DS2-GA29 complex is most stable at the global and regional level, KIR2DS2-GA20 complex is most stable at TYR45.

**Figure 12:**
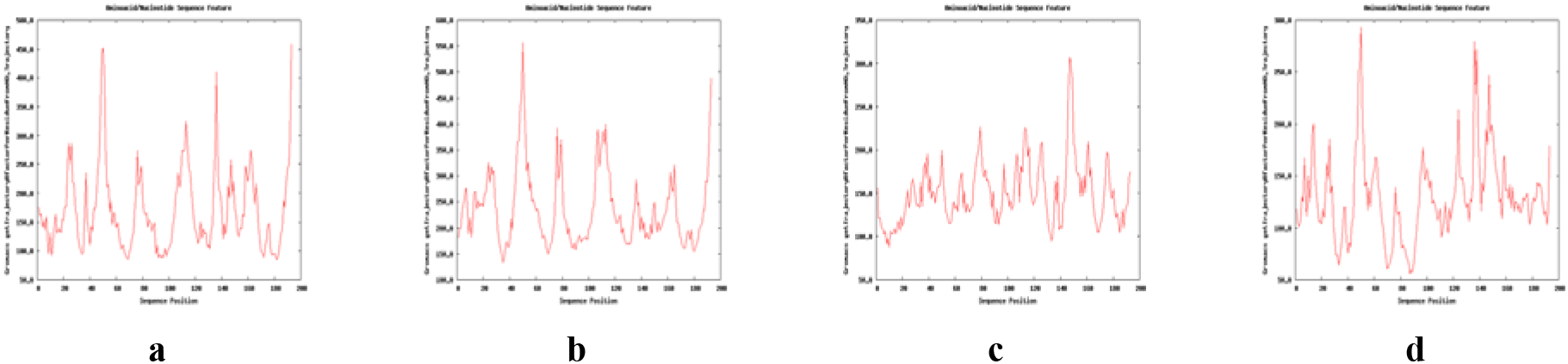
B Factor for Apo and Holo structures (a). KIR2DS2 (b) KIR2DS2-NAG (c) KIR2DS2-GA20 (d) KIR2DS2-GA29

### Principal components Analysis

Principal component analysis (PCA) can be used to determine the relationship between statistically meaningful conformations (major global motions) sampled during the trajectory [54]. From Figure 13 and Table 6, the global motions were least in the KIR2DS2 – GA29 complex. The KIR2DS2 – GA20 complex showed greater global motion than the KIR2DS2 – NAG complex. In a similar manner, the average motion of the first three principal components (PC1, PC2 & PC3) at residue 45 shows that the KIR2DS2 – GA29 complex is the most stable holo structure while the KIR2DS2 – NAG complex s more stable than the KIR2DS2-GA20 complex.

**Figure 13:**
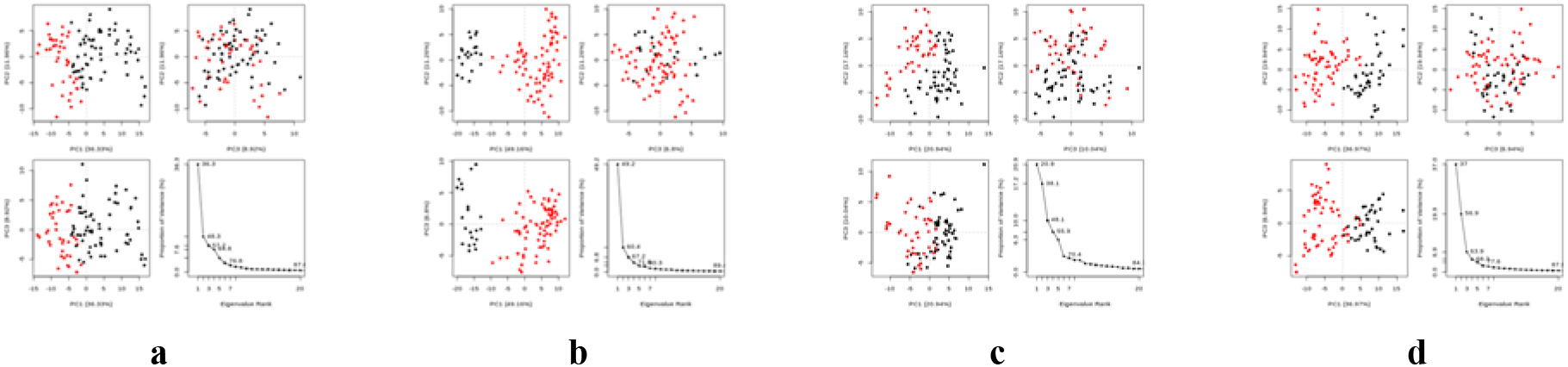
Principle component analysis cluster plot of Apo and Holo structures. The projection of trajectory onto 1st few eigenvectors for: (a). KIR2DS2 (b) KIR2DS2-NAG (c) KIR2DS2-GA20 (d) KIR2DS2-GA29

Based on the least motions, the best global conformations are PC1 of the Apo protein, PC2 of KIR2DS2 – NAG complex, PC3 of KIR2DS2-GA20 complex, and PC3 of the KIR2DS2 – GA29 complex. Similarly, the best conformations that produced the least motions at TYR 45 are PC1 and PC3 of the Apo protein, PC3 of KIR2DS2-NAG complex, PC3 of KIR2DS2-GA20 complex, and PC3 of the KIR2DS2–GA29 complex. The cosine contents of the principal components reveal the quality of the sampling and to what direction the MD simulation is converging (Table 6).

### Dynamic Cross Correlation Map (DCCM)

The DCCM reveals the heat map of cross correlation of residual fluctuations. The pairwise graph reveals the positive and negative correlation effects of the atomic displacements in the residues as they correlate with one another [55].

Figure 14 and Table 6, reveals a complex pattern of correlated, non-correlated and anticorrelated motions in Apo and Holo structures. Comparative results reveal that though more intensified, the atomic motions in the KIR2DS2-NAG complex resemble more closely that of the Apo structure. Similar to the Apo structure, approximately residues 1-50 (which includes TYR45) of the KIR2DS2-NAG complex shows correlated motion. The Ig-like c2 type 1 domain (42-107) of the same holo structure predominantly shows anticorrelated atomic motions. The map for Ig-like c2 type 2 domain (142-205) suggest that most residues predominantly showed non-correlated atoms. The heat maps of the KIR2DS2-GA20 and KIR2DS2-GA29 complexes closely resembled. They showed less anticorrelated and correlated motions, and more noncorrelated motions than the KIR2DS2-NAG complex in all regions of the protein.

**Fig 14:**
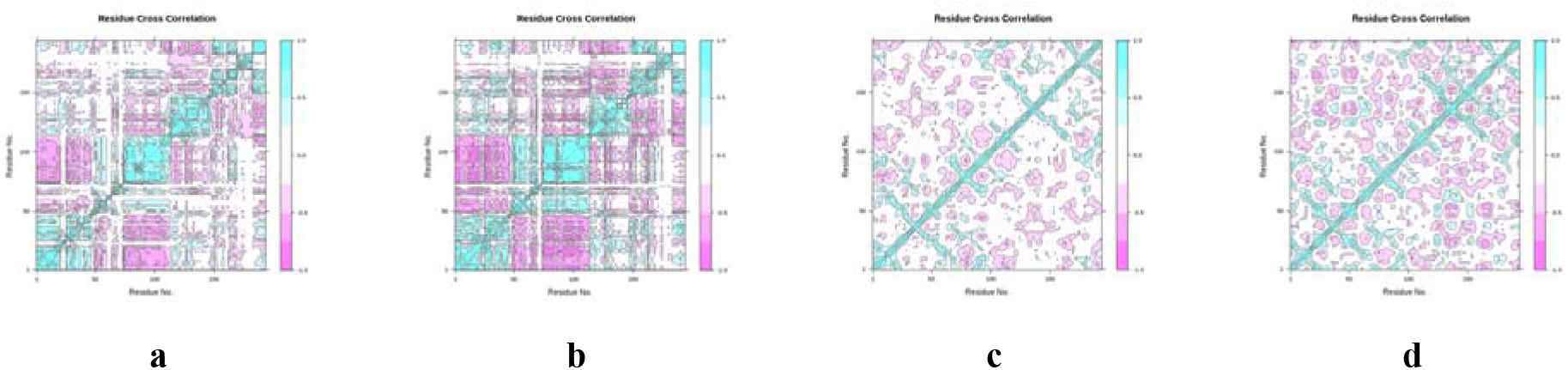
Dynamic cross correlation map Apo and Holo structures of 1m4k Purple represents anticorrelated, dark cyan represents fully correlated while white and cyan represents moderately and uncorrelated respectively. 1.0= correlated; 0 is non-correlated; and 1 is anti-correlated. (a). KIR2DS2 (b) KIR2DS2-NAG (c) KIR2DS2-GA20 (d) KIR2DS2-GA29.

## Conclusion

### The standard and lead compound

Glucosamine can be obtained from the shells of crabs, lobsters and shrimps. Orally administered NAG has been used in the treatment of Inflammatory Bowel Disease (IBD) and osteoarthritis [56]. The anti-cancer effect of NAG has also been well documented. By activating Death Receptor 5, NAG as an adjuvant promotes TRAIL-induced apotosis in Non-Small Cell Lung Cancer Cells [57]. Glucosamine also to exhibited antitumor activity in lung cancer cells by suppressing Forkhead box O proteins (FOXO) phosphorylation. Through a structural modification, Glucosamine specifically inhibited the translocation of FOX01 and FOX03 proteins which downstream signaling molecules for the PI3K/AKT and MAPK/ERK pathways [58]. Glucosamine has also been known to induce cell death in prostate cancer cells by the inhibition of proteasomal activity [59].

Gibberellins are plant hormones that regulate its various developmental processes and they are found abundantly in *Abelmoschus esculentus* (okra) and *Pisum sativum* (green peas) [60,61]. They are known to be involved in plant innate immunity by mostly by regulating SA–JA–ET signaling systems [62]. A Gibberellin derivative, GA-13315 has shown great promise as a chemotherapeutic agent against lung cancer [63]. Some gibberellin based molecules which show strong anticancer activities have also been designed and synthesized [64]. Specifically, the cytotoxic or immunomodulatory effect of GA20 and GA29 are yet to be ascertained.

### Summary of comparative analyses

Following a thorough evaluation of the bioavailability, pharmacokinetic properties and binding site analyses, NAG, GA20 and GA29 are predicted to be good drug candidates and have an immuno-stimulatory effect on KIR2DS2. However, GA29 would have the greatest immuno-stimulatory effect because it has the best binding affinity (−8.2 Kcal/mol) to the KIR2DS2 receptor and a greater nuclear receptor ligand and activity prediction than NAG. GA20 would also has a better binding affinity (−7.4 Kcal/mol) greater nuclear receptor ligand activity than the standard. Unlike, NAG, GA20 and GA29 are substrates of the CYP3A4 and this suggests that they should be administered with an inhibitor of that enzyme.

Studying the time-resolved motions of Apo and Holo macromolecules, GA20 and GA29 are predicted to have better pharmacodynamics on the target receptor than NAG. Specifically, of the two lead compounds, GA29 is predicted to have a better efficacy. This is because GA29 has shown better structural stability as seen with the values from the RMSD (values and distribution of peaks), RMSF (at TYR45), RoG (percentage gyration, range and average), B Factor (global and regional), and PCA (global and local motions). Furthermore, GA 29 has the strongest binding affinity to KIR2DS2 (−8.2 kcal/mol).

It is recommended that the immuno-stimulatory effect of Gibberellins A20 and A29 and NAG on KIR2DS2 be evaluated using *in vivo* and *in vitro* experiments.

